# Model of integrin binding as a function of light-induced variations in forces exerted on cells via RGD-terminated, photoswitchable azobenzene surfaces

**DOI:** 10.1101/2025.07.15.664514

**Authors:** Daniel A. Vaughan, Sophie Geiger, Anna M. Piccinini, Mischa Zelzer, Etienne Farcot, Christine Selhuber-Unkel, Bindi S. Brook

## Abstract

Photo-responsive biomaterials are attractive because of the ability to non-invasively alter and control the material properties, thus allowing control over the cell response at the interface with the bioamaterial surface. While in silico mathematical models have been implemented for simulating single-cell force spectroscopy (SCFS) experiments on static and some dynamic biomaterials, these models have yet to be extended to light-responsive biointerfaces. So here, we develop a mathematical model that describes and predicts the strength of integrin-mediated cell adhesions to a photoswitchable biomaterial surface. The fluctuating biomaterial comprises photoswitchable azobenzene attached to a glass surface and terminated with peptide c(RGDfK). Upon irradiation with light at 530 nm, the azobenzenes rapidly fluctuate between an extended and contracted conformation, leading to a change in the length of the azobenzene that stimulates integrins bound to c(RGDfK). The mathematical model mimics the nascent adhesion and spatial fluctuations in the extra-cellular matrix (ECM) and is calibrated using single-cell force microscopy retraction curves. It relies on spring-based mechanics to describe the stretch and deformation of a cell and ensembles of integrins when applied to a fluctuating biomaterial. We use the model to simulate retraction curves and the proportion of bound integrins on the surface as the material fluctuates. Additionally, we use the model to predict SCFS retraction curves for varying experimental conditions. This includes the length of time the cell (attached to the tip of the atomic force microscopy cantilever) is kept in contact with the biomaterial before retraction, and for varying frequency of light-induced movement of the azobenzene conformations. These model outcomes provide an attractive route to integrate and rigorously control certain experimental variables, thus accelerating experimental design of the study of cell adhesion to light-responsive biomaterials. Furthermore the model complements experiments by providing estimates of variables that are experimentally inaccessible. Thus, the outcomes of the model provide a valuable resource to aid in the interpretation and design of light-responsive biointerface functionalities.

## INTRODUCTION

Interactions between cells and the extracellular matrix (ECM) play an important role in cell survival, adhesion, proliferation, migration, and morphology [1]. These interactions are mediated through integrins that transduce mechanical forces from the ECM across the cell membrane and into the cell’s cytoskeleton [2, 3, 4, 5]. The transfer of force through integrins stimulates and activates mechanotransduction pathways [6, 7], which can play a pivotal role in tissue repair and neural regeneration [8, 9].

Approaches to probing mechanotransduction pathways activated by integrin stimulation [10] include single-cell force spectroscopy (SCFS) [11], optical and magnetic tweezers [12, 13, 14], and microscopy [15]. Among these, SCFS has previously been employed to quantify integrin-mediated adhesion forces between a single cell and a responsive surface that changes its cell-adhesive properties upon irradiation with light [11]. By linking the cell adhesion-promoting peptide – the cyclic RGD (arginylglycylaspartic acid) derivative c(RGDfk) – via a photoresponsive unit (azobenzene) to a glass surface, the availability of c(RGDfk) within the PEG environment is altered via photostimulation. This results in a change in the adhesion strength of fibroblast cells to this biomaterial surface that can be quantified via SCFS.

Modelling the interplay between cell-material adhesion strength enables understanding of the effect of different experimental parameters on integrin-mediated cell adhesion and allows exploration of variables that are not easily experimentally accessible. Current approaches include probabilistic evaluations (stochastic models) and evaluations of best parameter fits on continuous scales (continuum models) for cell shape and mechanotransduction-stimulated responses, such as Rac and Rho [16]. A continuum model approach has successfully modelled force transmission across the cell by defining local cellular motion as Hookean springs [17, 18]. The Motor-Clutch model [19] uses spring-based modelling to determine a cell’s optimum stiffness for maximal traction force. The authors provide an ordinary differential equation (ODE) model that adds validity to the Monte Carlo (MC) simulations for including motor-clutch dynamics in the model describing other areas of cell motility.

Integrins play a key role in affecting cell-adhesion strength to a biomaterial surface. Cell-adhesion strength is a variable that is experimentally accessible, e.g., via SCFS. In the past, several mathematical models have been developed to describe integrin-mediated cell adhesion strength to static biomaterials. To simulate the effect of loading on binding and unbinding of integrins between cell and ECM, Irons *et al*. have developed a continuum model combined with atomic force microscopy (AFM) data to quantify the proportion of bound integrins and adhesion strength between an ECM-functionalised AFM cantilever tip and the cell surface [20]. The simulations were compared qualitatively with the experimental AFM data to demonstrate their relevance and the ability to determine biologically relevant parameters. On the other hand, stochastic models, in which integrins are treated as individual springs, have also been implemented to quantify integrin binding and adhesion strength under loading forces [21, 22]. Additionally, [23] Erdmann *et al*. coupled stochastic simulations with dynamic force spectroscopy data to estimate the initial conditions for the simulation to predict maximum detachment forces. Although stochastic simulations can produce close representations of reality, parameter estimation can be computationally prohibitive, in contrast with continuum models. Furthermore, continuum models for integrin-responsive surface interactions have yet to be developed.

Most of these models have been developed to study cell interactions with static biomaterials, i.e., biomaterials that do not change their properties over time. Studies have incorporated aspects of dynamic changes in the biomaterial in their models [24], but no models exist that explicitly cater for the reversible dynamic changes in chemical properties induced by light on a photoresponsive bioamterial surface. The ability to use light as a non-invasive stimulus to change material properties in biological and biomedical applications is very attractive and intensely studied experimentally [25]. We therefore aim to fill this gap by developing a mathematical model for integrin-mediated force transduction that can be applied to dynamically changing, light-responsive biomaterial surfaces.

Here, we extend the model of Irons *e*t al. [20] to develop a mechanical model that couples integrin binding and photoswitchable molecule dynamics and simulates adhesion and detachment of a fibroblast from a surface using a cell attached to an AFM cantilever, to and from a photoswitchable azobenzene surface. Model parameters are inferred by fitting the model to SCFS retraction curves from work previously published by Kadem *e*t al. [26]. The resulting parametrised continuum model represents the interaction between fibroblast cells and photoswitchable biomaterials. This allows for predicting responses to new stimuli exposure regimes, facilitating the design of new materials and testing conditions that may ultimately lead to new biomaterial applications.

## METHODS

### Experimental Data

For the model in our work, previously published data from a study by Kadem *e*t al. [26] is used. In their SCFS experiments, Kadem *et al*. used a bioamaterial surface composed of a photoswitchable azobenzene attached to the glass substrate [26] (Figure 1b). c(RGDfk) was attached at the exposed end of the azobenzene, allowing for integrin-mediated cell attachment to the surface. The surface is populated with a 1:100 mixture of c(RGDfk)-terminated azobenzenes and polyethylene glycol (PEG 2000).

**Figure 1.**
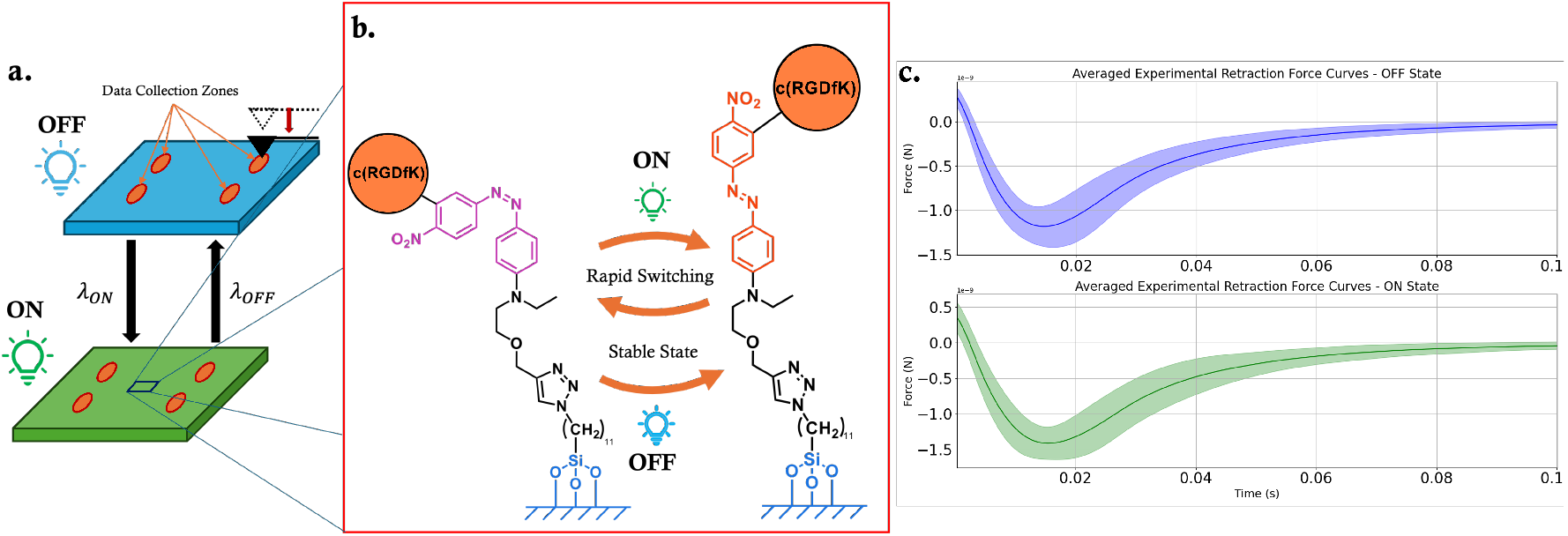
Experimental setup and data generated by single-cell force spectroscopy (SCFS). (a) The cell attached to an AFM cantilever is brought into contact with the photoswitchable c(RGDfk) terminated surface. Data is collected from four separate regions while the surface is either in the **OFF** or **ON** state. (b) Schematic of the photoinduced isomerism of surface immobilised azobenzene. In the **ON** state (*λ*_*ON*_ = 530 nm), azobenzene isomerism is in a dynamic equilibrium where azobenzene switches randomly between the *cis* and *trans* configuration, with a larger population in the cis state [[28] and [29]]. In the **OFF** state (absence of light), the azobenzenes exist in a steady state with a greater proportion in the extended state (*e, trans*) over the contracted state (*c, cis*). (c) The experimental force curve data from Kadem et al. [26] is averaged, producing the smooth retraction curves with shaded region depicting ±1 standard deviation (±*σ*), included to demonstrate the range of retraction curves collected.

Azobenzene exists in two isomers, a bent *cis* state (denoted *c* in the model) and a stretched *trans* state (denoted *e*). It can rapidly transition between these two isomers. While the extended *trans* state is thermodynamically more stable [27], both the *cis* and *trans* states coexist under **ON** lighting. In the absence of light (**OFF**-state), *trans*-azobenzenes are predominant. Upon irradiation with light at 530 nm (**ON** state), the prevalence of the *cis*-azobenzenes increases (Figure 1b).

For the SCFS experiment, a (REF52-WT) fibroblast cell adheres. The probe with the attached cell is brought in contact with the surface, allowed to remain in contact for a predetermined time (*T*_*p*_, either 1 s or 3 s) and then retracted from the surface. As the probe moves towards or away from the photoswitchable surface, the force exerted onto the cantilever is determined by measuring the deflection of a cantilever of known stiffness.

Approximately 20 SCFS force curves were collected at each of 4 separate locations on the surface (location 1–4) [26] (Figure 1a). The detachment force from every force curve was measured each time after switching the state of the surface, following the switching sequence OFF-ON-OFF-ON-OFF, such that data for the **OFF** state (3 times) and the **ON** state (2 times) is obtained for the same cell in the same location. Although collected sequentially, each curve is treated as a separate event here. Therefore, the completed dataset for the present model includes 794 curves, with 398 curves for the **ON** measurement and 396 for the **OFF** state. Figure 1c shows the resulting averaged data.

### The Model

The mathematical model for cell adhesion to surfaces undergoing light-induced changes in c(RGDfK) presentation is shown in Figure 2. At the microscale, the model comprises a system of coupled advection-reaction partial differential equations (PDEs), describing integrin binding and unbinding between the cell and cantilever, and the cell and the light-responsive biomaterial. We arrive at this by extending the model described by Irons *et al*. [20] by accounting for additional bound integrin complexes (ie those bound to contracted and extended azobenzene molecules, as well a generalising it to multiple layers (see supplemental material). The cell, probe attachment peptides, and individual integrins are treated as Hookean springs, and the rates of integrin binding and unbinding depend on the deformed lengths *L*_2_ (*µ* m) and *L*_4_ (*µ* m); this couples the microscale binding/unbinding to the macroscale deformations of the multipl layers during approach and retraction of the cantilever.

**Figure 2.**
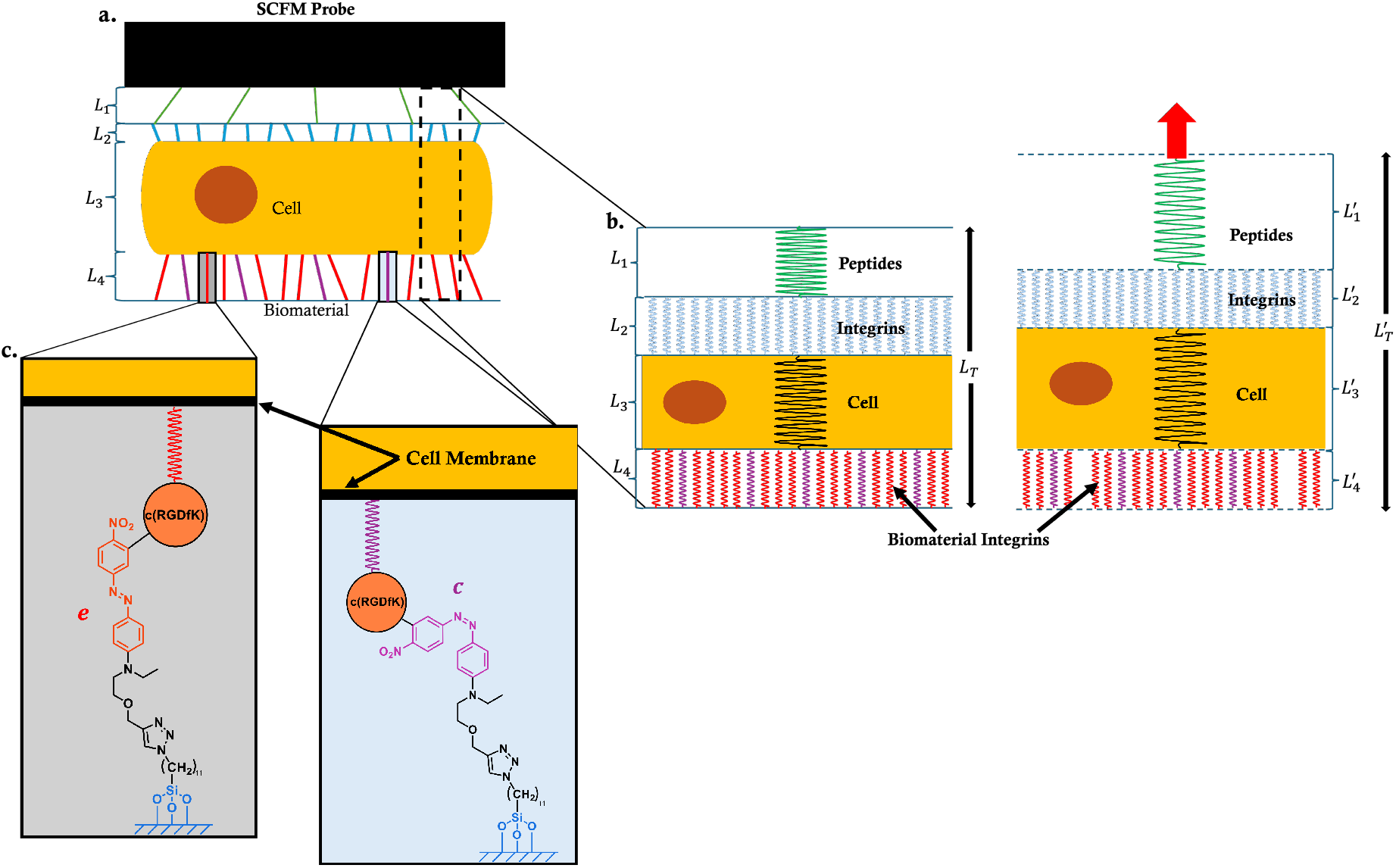
Diagram showing the abstraction of the experimental setup (a) and its conceptual translation into a mathematical model (b). Discrete layers with lengths *L*_*i*_, *i* = 1..4 are identified in the experimental setup: *L*_1_ is the length of the protein layer binding the cell to the tip of the cantilever; *L*_2_ is the length of the integrin layer connecting the cell to the protein layer on the cantilever; *L*_3_ represents the cell thickness; and *L*_4_ is the length of the integrin layer between the cell and the biomaterial surface. A spring model represents each layer, and the sum of the lengths is *L*_*T*_. Subscripts *L*_*i*_, *i* = 1..4 represent the natural length of the springs under zero loading (b, left), while subscripts 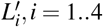 represent deformed lengths of the layers that vary with time (b, right). The upward movement of the cantilever during retraction is depicted in (b, right); during the approach, the springs would be compressed. In layer 4, the red and purple colours indicate that integrins bound to azobenzene can exist in either the extended *trans* (*e*) or contracted *cis*(*c*) state, respectively, and these proportions also vary with time. (c) The switching of the photo-sensitive azobenzene from extended *trans* (*e*) to contracted *cis*(*c*) and illustrating the deformation in spring length. The spring represents the integrin attached from the cell membrane to the c(RGDfk).

### Integrin dynamics

The equations governing the dynamics of integrins in the layer connecting the cell and the biomaterial (*L*_4_ in Figure 2) are represented by:

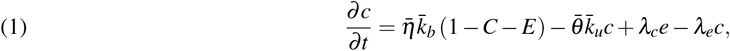

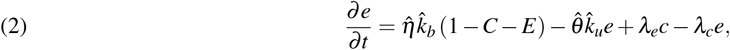

*c*(*x, t*) and *e*(*x, t*) are the proportion of integrins bound to the contracted or extended azobenzene molecules, respectively. Each bound integrin is associated with a microscale coordinate or horizontal displacement *x, local* to each integrin, that indicates the distance from its resting/undeformed position to a c(RGDfK) binding site (Fig 2c). Therefore, the continuous variables *c* and *e* are, in fact, distributions as functions of *x* that can vary with time *t. C*(*t*) and *E*(*t*) are the total proportions of integrins bound to contracted and extended azobenzene molecules, respectively, given by

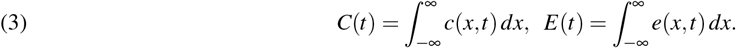

Since these are proportions of the total integrins available for binding, we assume that at any time *t, C* + *E* +*U* = 1, such that the quantity *U* = 1 −*C* −*E* in equations (1) and (2) is the total proportion of unbound integrins *U* (*t*), and 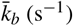 and 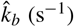 are the binding rates of unbound integrins to either the contracted or extended azobenzene molecules, respectively. The parameters 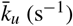 and 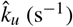 represent the unbinding rates of integrins bound to contracted and extended molecules, respectively. These binding and unbinding rates depend on the extension of the integrins and are described in more detail below. The parameter *λ*_*c*_ represents the rate at which integrins bind to azobenzenes in the contracted *cis* state and switch to the extended *trans* state, and *λ*_*e*_ (s^−1^) represents the rate of the opposite reaction; these are light-dependent and detailed below. 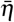, and 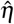 are dimensionless parameters which modulate 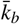 and 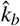 respectively and are factors reflecting the availability of binding sites on the underside of the cell with the available c(RGDfk). 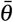, and 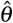 are dimensionless parameters which modulate 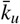 and 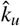, respectively, and determine the relative stability of the bond formed by the integrin. Following Irons *et al*. [20], the equation that governs the dynamics of bound integrins *b*(*x, t*), between the cell and probe (layer 2 in Figure 2), is given by,

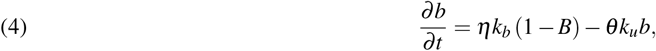

where *k*_*b*_ (s^−1^) and *k*_*u*_ (s^−1^) are the rates of binding and unbinding of integrins to proteins on the SFCS probe, respectively, and *B*(*t*) is the total proportion of bound integrins in this layer and given by

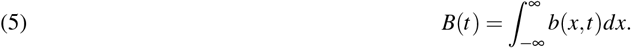

Therefore, 1 −*B* is the total proportion of unbound integrins available for binding. *η* and *θ* are dimensionless parameters which modulate the binding and unbinding rates, respectively; the rationale for their inclusion is the same as for the scale factors (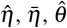, and 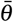) that appear in equations (1) and (2).

The binding and unbinding rates are modelled as piecewise linear functions of integrin extension during retraction, 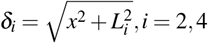 (e.g. Fig. 2d), such that

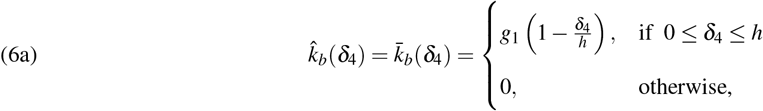

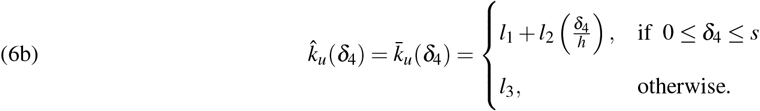

Similarly

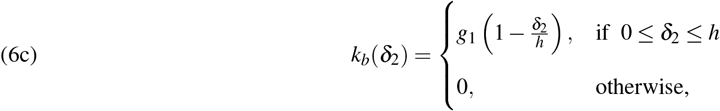

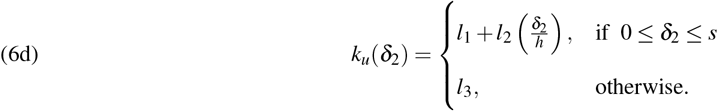

Since *L*_2_(*t*) and *L*_4_(*t*) will be determined as part of the mechanical model (see below), and hence known at each time point, *δ*_2_ and *δ*_4_ are effectively functions of *x*. The parameters *h* and *s* indicate integrin extensions above which binding rates are zero and unbinding rates are at their highest value, respectively. The parameter *g*_1_ (s^−1^) is the maximum binding rate, and *l*_1_ (s^−1^) is the unbinding rate when the integrin rests. The unbinding rate increases with linear gradient *l*_2_*/h* (s^−1^) as the integrin extension (*δ*_*i*_) increases. After a maximum extension *δ*_*i*_ = *s*, any remaining bound integrins are forced to unbind with a high rate *l*_3_ (s^−1^). These definitions and corresponding values are summarised in Table 1 in the supplementary).

**Table 1.**
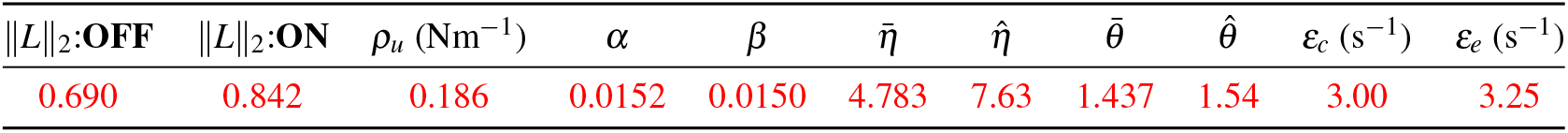
Fitted parameters from the OFF and ON datasets. Presented with their respective ∥*L*_2_∥ values.

The rates *λ*_*c*_ s^−1^ and *λ*_*e*_ s^−1^ capture the switching between *cis* (contracted) and *trans* (extended) states of the azobenzene molecules. They are modelled as piecewise constant functions of light intensity. Since we only consider light **ON** and light **OFF** states, we assume that when the light intensity *ξ* > 0, the light is in the **ON** state and **OFF** otherwise. Hence, the rates can be modelled via

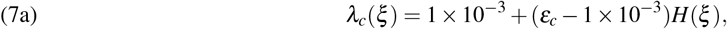

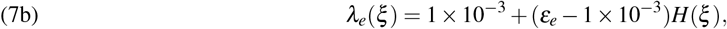

where *H*(*·*) is the Heaviside function. Thus, when *ξ* < 0 (in the light **OFF** state), *λ*_*c*_ = 1 × 10^−3^ s^−1^ and *λ*_*e*_ = 1 × 10^−3^ s^−1^ ensuring that in the OFF steady state, no switching is occurring as illustrated in Figure 1. The values selected for *ε*_*c*_ and *ε*_*e*_ are an assumption. Alternatively, when *ξ* > 0 (in the light **ON** state), the value of *λ*_*c*_ = *ε*_*c*_ > 1 × 10^3^ and the value of *λ*_*e*_ = *ε*_*e*_ > 1 × 10^−3^ ensuring that there is switching present and approximately equal proportions of integrins are bound to azobenzenes in the extended *trans* and contracted *cis* states. In reality, there is rapid switching between the two states in the light **ON** state (Fig 1); because this is so rapid, we assume that *ε*_*c*_ and *ε*_*e*_ can be considered to be constants on the observable timescale and thus exist in a dynamic equilibrium. [30].

### Mechanical model of the system

Each layer is modelled as a spring such that the system is considered a set of springs in series. Hence, the force (or, specifically, the tension) in each spring is given by

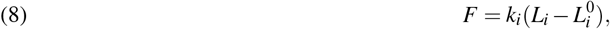

where for each layer *i, k*_*i*_ (Nm^−1^) is the spring constant, 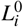 is the undeformed spring length (under zero loading), and *L*_*i*_(*t*) is the deformed length of the spring. Layers 1 and 3 are treated as Hookean springs; therefore, the spring constants *k*_1_ (Nm^−1^) and *k*_3_ (Nm^−1^) are constants. In layers 2 and 4, bound integrins are treated as individual Hookean springs in parallel; however, because the total proportion of bound integrins (*C, E, B*) vary with integrin extension, *δ*, the effective spring constants *k*_2_ and *k*_4_ are nonlinear and given by

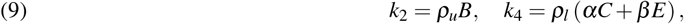

where *ρ*_*u*_ and *ρ*_*l*_ are factors that are given by multiplying the spring constant of individual integrins with the proportion of integrins bound on the upper and lower cell surfaces. The parameters *α* and *β* represent the contributions to *k*_4_ of the integrin-bound azobenzene molecules in the contracted and extended states.

The model has been built to accommodate any number of layers adaptable to different systems. At any time *t*, the force balance equations for *N* springs in series representing *N* layers can be written as:

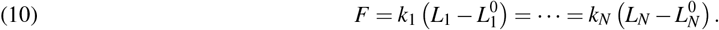

In our present experimental setup (Fig 2b), we assume that in layer 2, the total proportion of bound integrins 0 < *B* ≤ 1 throughout retraction (i.e., *B≠* 0), based on the experimental observation that the cell remains bound to the cell attached to an AFM cantilever tip during the experiment. On the other hand, in layer 4, we assume the proportions of integrins bound to the azobenzene molecules 0 ≤ *E* ≤ 1 and 0 ≤ *C* ≤ 1, allowing total detachment of the cell from the biomaterial. At this point, *E* = *C* = 0 and layer 4 is decoupled from the rest of the layers. Consequently, we have to consider two separate cases for the state of Layer 4:

**Case (i)**: Either (0 ≤ *E* ≤ 1 and 0 < *C* ≤ 1) OR (0 < *E* ≤ 1 and 0 ≤ *C* ≤ 1). In this case, some integrins are bound to extended or contracted azobenzene molecules, so all four layers are coupled such that *i* = 1..4. Therefore with *N* = 4, the force balance equations (10) become:

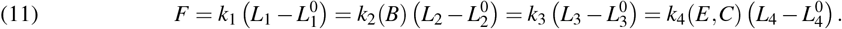

The total length of the system in the absence of loading is defined by

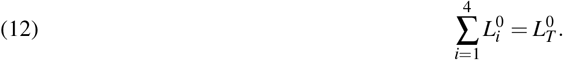

The spring system is driven by the displacement of the cell attached to an AFM cantilever, *L*_*T*_ (*t*), which is prescribed based on the experimental protocol and given by

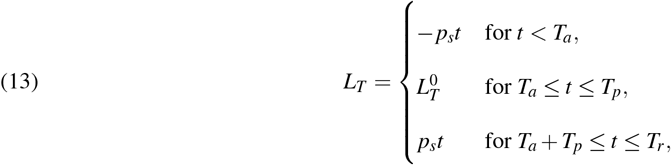

where *p*_*s*_ > 0 is the speed of the probe as defined in [26] to be *p*_*s*_ = 30 *µ*m/s, *T*_*a*_ is the time point at which the cell first touches the biomaterial surface, *T*_*a*_ + *T*_*p*_ is the time point at which retraction of the probe begins, and *T*_*r*_ is the time point at which the probe stops retracting. *T*_*a*_, *T*_*p*_, and *T*_*r*_ are defined experimentally and will have slightly varying values from one retraction curve to another. However, an averaged value for *T*_*a*_ = *T*_*r*_ = 0.34 s, the distance that the probe will travel should be enough to ensure complete detachment of the cell from the biomaterial as was achieved in the experiment.

Since the individual layers are connected at any time *t*, it is also true that

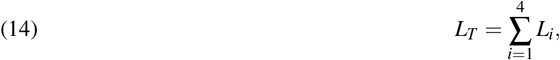

By sequentially eliminating the unknowns from equations (11) – (14) (see Supplementary Material), we obtain the following generalised equation for the length of the *i*th layer at any time *t*:

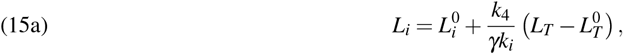

where

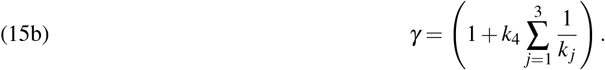

**Case (ii):** *E* = 0 and *C* = 0. In this case, the cell detaches completely from the biomaterial surface. Therefore, layer 4 is no longer connected to the other layers, and the system becomes unloaded such that *F* = 0; equation (11) thus immediately yields 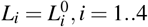, *i* = 1..4. Hence, as expected, each layer returns to its resting length upon unloading.

### Computational simulations

Equations (1) – (5), (11) and (15) form a coupled system of equations for Case (i). These are solved numerically for the variables *c*(*x, t*), *e*(*x, t*), *b*(*x, t*),*C*(*t*), *E*(*t*), *B*(*t*), *F*(*t*) and *L*_*i*_(*t*), *i* = 1..4 subject to the initial conditions:

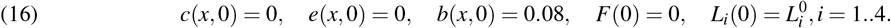

The boundary conditions *c* → 0, *e* → 0, and *b* → 0 as *x* → ∞ are automatically satisfied through the high unbinding rates imposed. The initial condition for the layers are *L*_1_(0) = 100 nm, *L*_2_(0) = 10 nm, *L*_3_(0) = 200 nm, *L*_4_(0) = *L*_*T*_ (0) − (*L*_1_(0) + *L*_2_(0) + *L*_3_(0)). These values are the same as what was used by Irons *et al*. in [20].

### Parameter Fitting

As shown in Fig. (1), in the experiments, force is measured as a function of time during the approach and then retraction of the cell attached to an AFM cantilever. Using the coupled model detailed above, *F*(*t*) can be simulated for a range of parameters and compared directly with these experimental data in both light **ON** and **OFF** states. The parameters obtained from the literature are fixed. However, not all parameters could be obtained in this way. We, therefore, first estimate the set of model parameters 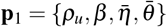 by fitting the model to the SCFS retraction curves in the **OFF** state: Once these parameters have been estimated, the parameters 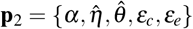 are estimated by fitting the model to the SCFS retraction curves in the **ON** state, since these parameters are specific to simulating the **ON** state.

The parameter estimation is performed by evaluating the ∥*L*_2_∥, given by,

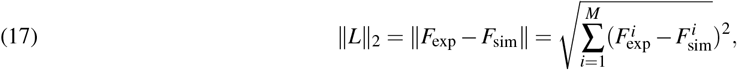

where *M* is the number of data points in the averaged SCFS retraction curves. The minimisation of the objective function represented in equation (17) produces the set of parameters that can be used in further simulations. The values from *F*_sim_ required to perform the minimisation are selected from the same time points as the experimental force data point.

The 50 initial parameter sets of *p*_*i*,1_, *i* = 1, …, 50 were generated using the Latin Hypercube Sampling (LHS) method. The ranges used for generating the initial estimation for *p*_*i*,1_ were found from a coarse exploration of the parameter space to identify a region that provided a low ∥*L*∥_2_. After the optimization for the *p*_*i*,1_ sets, a further set of parameters is generated for *p* _*j*,2_, *j* = 1, …, 250. Every 5 sets of *p* _*j*,2_ are paired with a set from *p*_*i*,1_ and optimised on the **ON** retraction curves until the lowest possible ∥*L*∥_2_ is reached. The parameters are selected based on the suitability of *p*_*i*,1_ and the paired *p* _*j*,2_. This process is beneficial as the number of parameters fitted to the **OFF** and **ON** retraction curve is reduced.

## RESULTS

The selected parameters (**p**_1_ and **p**_2_) from the parameter estimation are used to investigate how different experimental conditions (by altering the parameters in the model) affect the retraction curve, thus providing predictions that can inform and be tested in future experiments. To observe the influence of selected parameters on the resulting SCFS retraction curve, we investigate variation in the following conditions:

i. contact duration (*T*_*p*_) between the cell and the biomaterial surface;
ii. speed at which the probe (*p*_*s*_) is retracted from the surface of the material; and
iii. frequency of switching the light (*f*) **OFF***↔***ON**;

These conditions have been selected due to the ease of changing in the experimental setup (varying *T*_*p*_ and *p*_*s*_) and for results that would be unobtainable without significant variation in the experimental setup (varying *f*). From this analysis, further experiments are proposed for additional information to improve the robustness of the model and increase understanding of the system.

### Error Analysis and Selection of Parameter Sets

Parameters are selected based on their respective ability to produce simulations that closely resemble the averaged experimental retraction curve (for both **OFF** or **ON**). The accuracy of the simulation is assessed through the ∥*L*∥_2_ value and visual overlay of the simulations and experimental data. Another parameter constraint is the proportion of integrin binding for *E* and *C*. Experimental data from [26] indicates that in the **OFF** state, the proportion of integrins bound to the extended azobenzene (*E*) should be significantly higher than that of the contracted azobenzene (*C*). In addition, it was shown that the proportion of integrin binding in the **ON** state would, at any given time, lead to an equal proportion of integrin bound to extended azobenzene (*E*) and contracted azobenzene (*C*). The extensive simulated datasets for **p**_1_ and **p**_2_ are provided on GitHub along with the code required to perform the model optimisation.

### Selected Parameters

Following the parameter estimation process outlined in the methods section, the parameters shown in Table 1 were selected to investigate potential experimental conditions computationally. A comparison of the simulated and experimental retraction curves for the optimised parameter set in Table 1 is shown in Figure 3. The simulated curve for the **OFF** and **ON** state is within ±*σ* (±1 standard deviation) of the averaged experimental curve.

**Figure 3.**
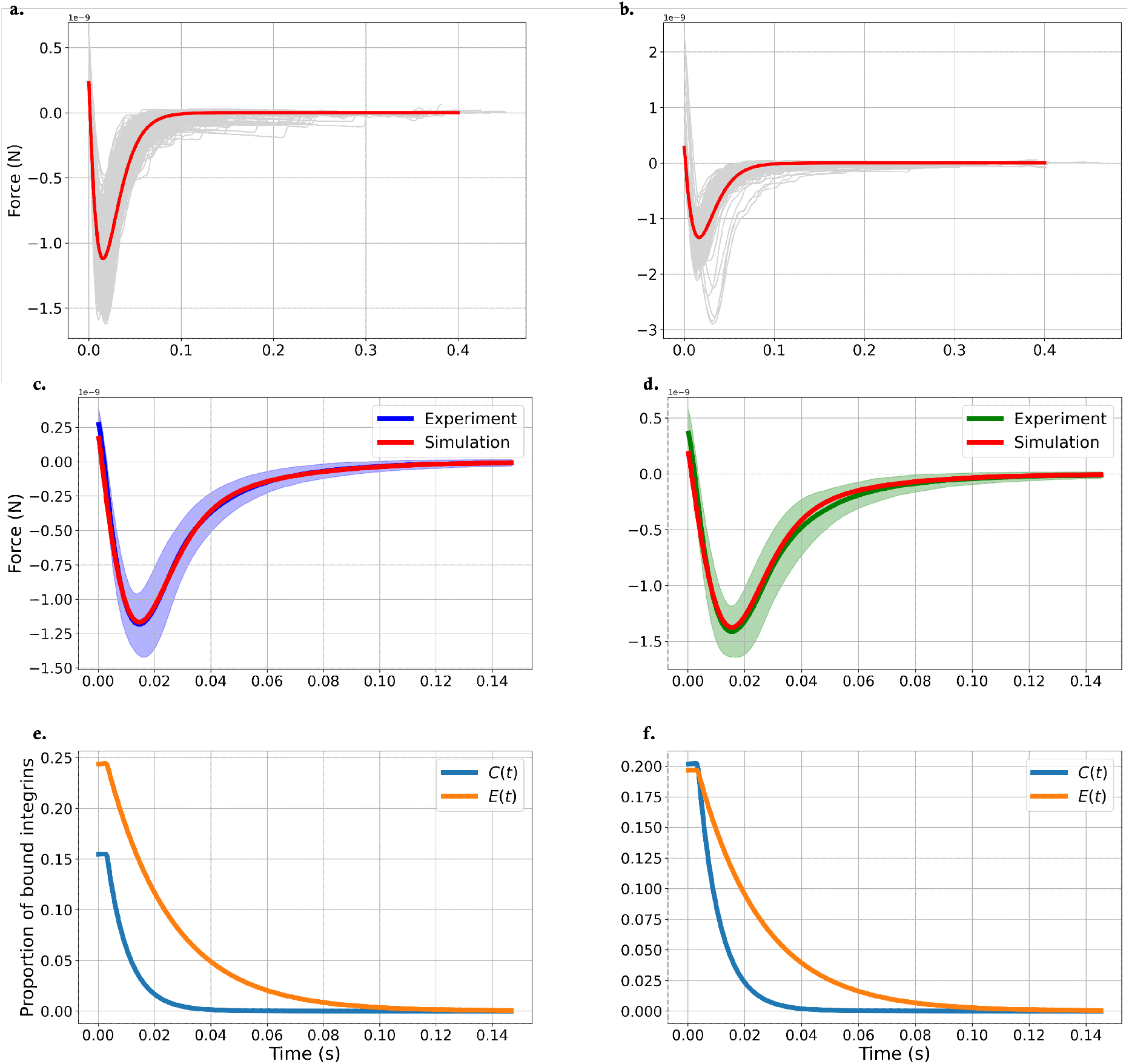
Simulated and average experimental force retraction curves in the **OFF** (a, c) and **ON** (b, d) states and simulated proportion of bound integrins in both states (e, f).In (a) and (b), simulated and all experimental retraction curves (light grey lines) are plotted together, illustrating the similarity between the simulations (red lines) after the parameter estimation and the full experimental dataset. In (c) and (d), experimental data (dark purple and dark green lines) is shown with one standard deviation (±*σ*, light purple and light green areas). Simulated retraction curves (red lines) were generated using the optimised parameter set (Table 1). (e) and (f): Total proportion (*C, E*) of integrin bound to c(RGDfK) terminated azobenzene surfaces in the **OFF** (e) and **ON** (f) state as the cell attached to an AFM cantilever is retracted over time.

### Estimation of the proportion of bound integrins

The simulation includes variables that describe the proportion of integrins bound to c(RGDfK) on either extended (*trans*-isomer) (*e*) or contracted (*cis*-isomer) (*c*) azobenzene molecules. This data is not accessible experimentally and presents a key benefit of modelling. The total proportion of integrin bound to c(RGDfK)-*trans*-azobenzene (*E*) and integrin bound to c(RGDfK)-*cis*-azobenzene (*C*) throughout the retraction is shown in Figures 3c and d.

In Figure 3, *E* and *C* show clear differences in the proportion of integrins bound to c(RGDfK) on *trans*-azobenzene and *cis*-azobenzenes on surfaces in the **OFF** and **ON** state. In the **ON** state (Figure 3d), *E* > 0 and *C* > 0 at the start of the experiment (*t* = 0). In contrast, in the **OFF** state, a higher proportion of integrins are bound to c(RGDfK) on *trans*-azobenzene, *E* > 0, while *C* = 0 at *t* = 0 (Figure 3c). This data suggests that modulation of cell-surface adhesion strength as a result of light stimulation of the c(RGDfK) azobenzene surface is related to the change in the proportion of integrins connected to azobenzenes in the *trans* or *cis* state.

### Computational Experiments

To perform SCFS, the experimentalist has to set a series of experimental parameters. These include the contact time *T*_*p*_ of the cell on the SCFS tip with the biomaterial surface and the speed at which the cell attached to an AFM cantilever is retracted from the surface. The choice of parameter values for these two parameters is expected to affect the measurement outcome, i.e., the measured cell adhesion force is a function of *T*_*p*_ and the retraction speed, among other parameters. It is reasonable to assume that *T*_*p*_ affects the proportion of integrins bound and that the retraction speed affects the measured force. The light-responsive azobenzene system explored here includes additional levels of complexity. Even if the intensity of the light is kept constant, the experimentalist must choose the frequency and length of the stimulating light (530 nm) used to induce the **ON** state.

Due to the large number of experimental measurements required to obtain a robust dataset for a single set of conditions, it is challenging to explore the parameter space fully experimentally. In the following section, we use our model to examine the dependence of the force curve characteristics and resulting adhesion forces by varying one parameter at a time.

### Effect of increased contact time on adhesion force and proportion of bound integrins

Parameter fitting in previous sections was undertaken using a contact time of *T*_*p*_ = 1 s between the SCFS tip and the c(RGDfK) terminated azobenzene surface. This corresponds to the experimental conditions under which the data used for the fitting was acquired [26]. Here, we explore if the model can generate predictions for a longer contact time of *T*_*p*_ = 3 s.

A comparison of simulated force curves and predicted relative integrin binding to c(RGDfK) in either the **OFF** or **ON** state for *T*_*p*_ = 1 s and *T*_*p*_ = 3 s is shown in Figure 4a. The *T*_*p*_ = 3 s simulation shows a higher attachment force (Figure 4a.i) and higher relative numbers of integrin binding with extended and contracted c(RGDfK) terminated azobenzene molecules in both the **ON** and **OFF** state compared to the *T*_*p*_ = 1 s data. This matches the experimental observation that more force is required to detach the cells from the c(RGDfK) terminated azobenzene surface if *T*_*p*_ = 3 s [26].

**Figure 4.**
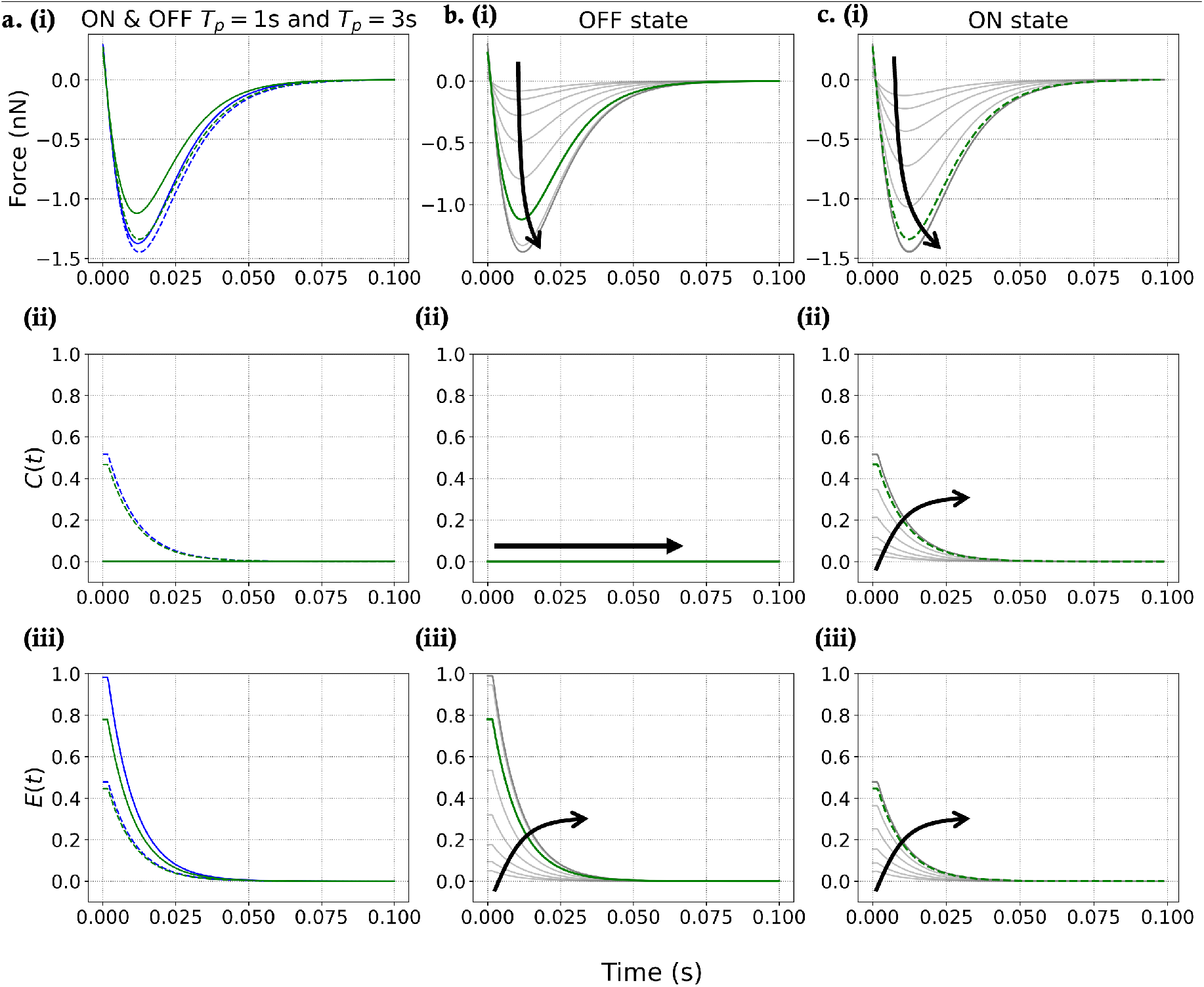
Simulated force curves (i) and relative amount of integrins bound to *cis*-azobenzene (*C*) (ii) and *trans*-azobenzene (*E*) (iii) for varying cell-surface contact times (*T*_*p*_) on surfaces in either the **OFF** or **ON** state. (a) Simulated data for the **OFF** (solid lines) and **ON** (dotted lines) state for *T*_*p*_ = 1 s (green) and *T*_*p*_ = 3 s (blue). (b,c) Simulated data for a range of contact times (*T*_*p*_ ≫1 and *T*_*p*_ ≪ 1) for surfaces in either the **OFF** (b) or **ON** (c) state. The (green) line represents *T*_*p*_ = 1 s; grey lines indicates *T*_*p*_ > 1 s and *T*_*p*_ < 1 s. Arrows indicate the effect of increasing *T*_*p*_. For all parameters other than *T*_*p*_, the optimised and predefined parameter values given in Tables 1 are used. For the **ON** state, *λ*_*c*_ = *ε*_*c*_ and *λ*_*e*_ = *ε*_*e*_. For the **OFF** state, *λ*_*c*_ = 1 × 10^−3^ and *λ*_*e*_ = 1 × 10^−3^.

The effect of a further increase and decrease in *T*_*p*_ is simulated in Figure 4b & c. The graphs in Figure 4b.i & c.(i) show an increase in the maximal retraction force - and hence an increase in cell adhesion force - as *T*_*p*_ increases. This agrees with simulations previously reported by Irons et al. [31]. For the same *T*_*p*_, the simulated attachment forces are always higher for the **ON** state than the **OFF** state, matching previous experimental observations of higher attachment forces for *T*_*p*_ = 3 *s* vs *T*_*p*_ = 1 *s* [26]. This trend in the simulation is accompanied by an increase in the relative number of bound integrins for all scenarios as *T*_*p*_ increases. We interpret this with the suggestion that longer *T*_*p*_ gives the integrins more time to form and mature before retraction of the SCFS tip begins. Based on the binding rates found in the optimised parameter set in Table 1, it can be seen that the detachment force, as well as *C* and *E*, reach a saturation point. This presents a prediction for the upper limit (−1.48 pN) in the retraction force and provides an estimate for *T*_*p*_ = 2 *s* by which the maximum binding has been attained.

### Effect of increased probe speed on adhesion force and proportion of bound integrins

The retraction speed of the AFM probe in the SCFS experiment may affect the measured adhesion force of the cell to the c(RGDfK) terminated azobenzene surface. The model incorporates the probe’s speed through the parameter *p*_*s*_ as shown in equation (13) and assumes that the cantilever’s acceleration is zero in the approach and the retraction. The simulations presented so far have been undertaken at a retraction speed of 30 *µ*m/s to match the instrumental parameters used to acquire the experimental dataset [26]. The simulated curves in Figure 5 show how different retraction velocities affect the model output. In these simulations, the travel distance of the cantilever was kept constant. Hence, the time required for the cantilever to retract from the surface varies for each simulation.

**Figure 5.**
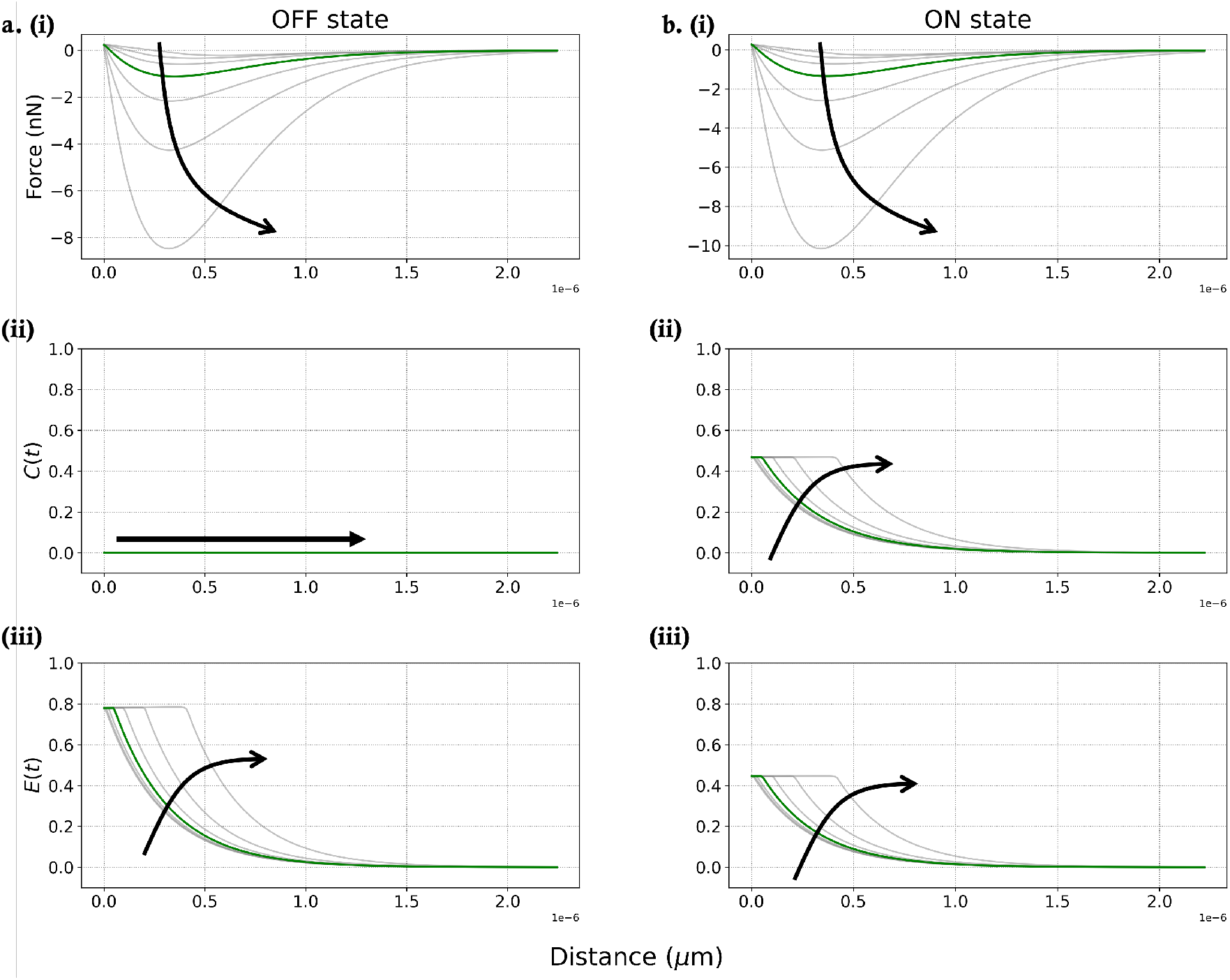
Simulated force curves (i) and relative amount of integrins bound to *cis*-azobenzene (*C*) (ii) and *trans*-azobenzene (*E*) (iii) for varying retraction speed of the cell attached to an AFM cantilever on surfaces in either the **OFF** (a) or **ON** (b) state. The (green) line represents the speed used in Reference [26] (30 *µ*m/s); data from other retraction speeds are shown in grey. Arrows indicate the effect of increasing speed. The parameters in Table 1 were used for all simulations. For the **ON** state, *λ*_*c*_ = *ε*_*c*_ and *λ*_*e*_ = *ε*_*e*_. For the **OFF** state, *λ*_*c*_ = 1 × 10^−3^ and *λ*_*e*_ = 1 × 10^−3^.

The simulated retraction force increases with the increase in the retraction speed of the cantilever. For low speeds, the probe does not leave the region of exponential unbinding until later. This causes an increase in the amount of binding, followed by a delayed exponential rate of unbinding (Figure 5a, b ii and iii). Based on this data, we suggest that reducing shock-stress events through rapid retraction allows integrin binding to be maintained for longer. In addition, the slower retraction speed allows more time for molecular rearrangements to take place, thus reducing the force required to detach the cell.

### Biomaterial Fluctuations During Adhesion

A key new feature of this model is the ability to simulate the effect of the external (photo-)stimulus on the proportion of bound integrins. The model enables changes of the proportions of *cis*- and *trans*-azobenzenes – which is synonymous to the photo-induction of the **ON** and **OFF** states in the model – over time and creates simulated retraction curves for these scenarios. Using this approach, we can connect different photo-stimulation patterns of the system and the resulting integrin binding state to the cell detachment force.

### Periodic Light Switching

Switching the light on the photosensitive biomaterial periodically creates a sinusoidal pattern in the proportion of integrins bound to the cell and the biomaterial surface (*L*_4_). For the periodic switching, the two factors that could affect max |*F*| are the frequency of how often the light switches between **OFF** and **ON** and the length of contact time (*T*_*p*_).

In this simulation, the light is varied by switching the parameters *λ*_*c*_ = 1 × 10^−3^ → *ε*_*c*_ and *λ*_*e*_ = 1 × 10^−3^ → *ε*_*e*_ for the **OFF**→**ON** transition. Switching happens regularly at a desired frequency (*f*). From Figure 6(b, c), a sinusoidal patterns appear for the proportion of total bound integrins (*E, C*) with the cell and the azobenzenes in the *cis*s or *trans* isomerism form from the switching driven by the light. This creates two components to analyse: the frequency and the phase for the sinusoidal pattern that forms in the proportion of total integrin binding in the biomaterial layer.

**Figure 6.**
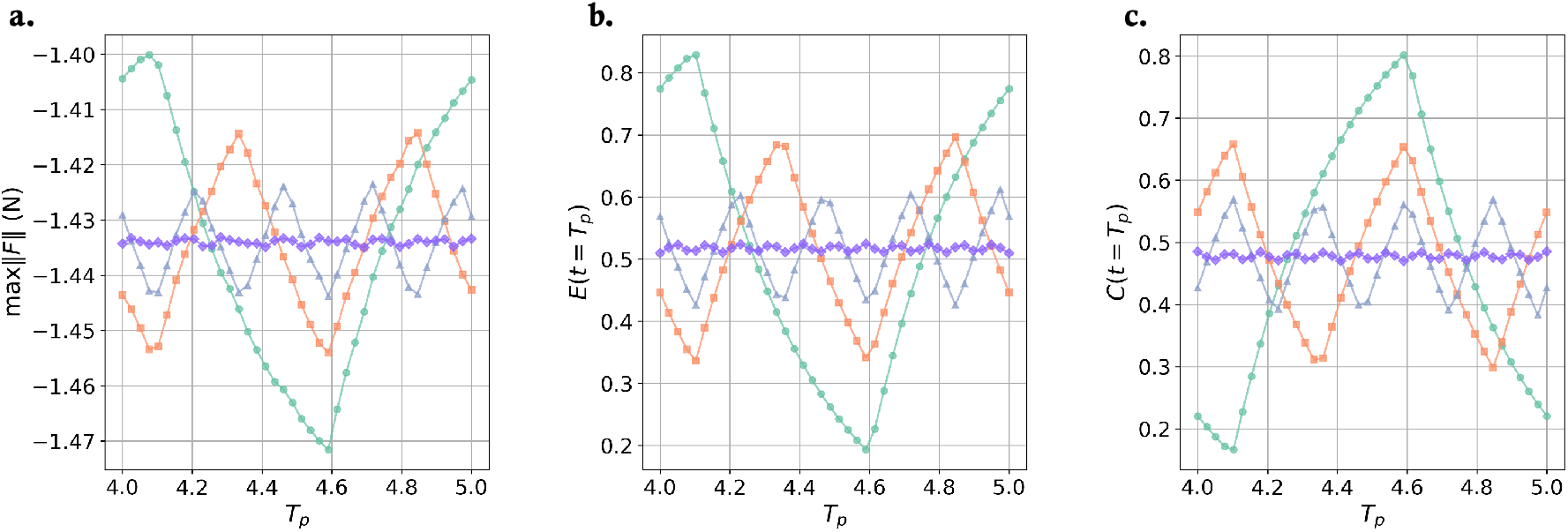
Effect of varying frequency (*f* = 2.0, 4.0, 8.0, and 100.0) of light stimulation on the simulated relative integrin binding and detachment forces. (a) Maximum detachment force experienced while *T*_*p*_ varies for the various switching frequencies. (b, c) Effect of contact time (*T*_*p*_) on the proportion of total integrin binding to c(RGDfK) terminated *trans*- and *cis*-azobenzene for the switching frequencies at the point when retraction is initiated. As the length of *T*_*p*_ increases, the retraction happens later, causing the sinusoidal nature of the integrin binding when the light is switched at a regular frequency.

The frequency of the exposure to light affects the proportion of integrins bound to c(RGDfK)-terminated (trans)-azobenzene (*E*) and (cis)-azobenzene (*C*) (Figure 6a). From Figure 6, it can be seen that for the lower frequency (*f* = 2.0 s^−1^), the *F*_max_ ranges from −1.40 pN to −1.471 pN. While this is a noticeable difference in the computational simulations, this would not appear outside the region of ±*σ* for experimental data, making the observed difference difficult to quantify experimentally. For higher frequencies *f* = 8.0 s^−1^, the range of *F*_max_ reduces further and ranges between −1.427 pN to −1.443 pN. Higher frequencies (*f* = 100.0 s^−1^) further decrease the amplitude of the detachment force. Therefore, a noticeable trend indicates that increased switching frequency reduces the amplitude of the oscillatory adhesion force.

## CONCLUSIONS

Stimuli responsive biomaterials are widely explored to exert dynamic control over cell-material interactions. Light-responsive materials are of particular interest because of the non-invasive nature of the photostimulus. While previous mathematical models have described integrin-mediated force transduction in static and dynamic materials, no mathematical models have yet been reported that incorporate light-induced changes of material properties and predict integrin-mediated force trasnduction in such systems.

To develop a mathematical model of integrin-mediated force transduction that takes light-induced, dynamic material changes into account, we have adapted a previous integrin binding model to simulate single cell force spectroscopy (SCFS) measurements based on treating the cell-integrin-material system as multiple connected spring layers adapted in this work to include characteristics of a responsive biomaterial. In our model, we account for the dynamic nature of a photoswitchable azobenzene surface by including variables that represent the extended, *trans*-azobenzenes and contracted, *cis*-azobenzenes bound to integrins via c(RGDfK) functionalities. In addition, the force balance equation has also been generalised to handle any *N* layers in the system, enabling the construction of models with a more complex system in the future.

The model has been calibrated with experimental data from SCFS retraction curves previously published in [26]. The parameter fitting optimisation provided a range of optimal parameters that produce a simulated retraction curve that recapitulates the experimental data in ±*σ* of an averaged retraction curve.

The computational simulations thus allow for the analysis of other aspects of the system with a degree of confidence. We investigated the following experimental conditions computationally: variations in the contact time between the cell and the surface, different speeds of the probe during retraction, and fluctuations in the azobenzene isomer composition to mimic the effect of photostimulation.

Detachment force curves, for 1- and 3-second contact times, agreed well with the data from Kadel et al’s experimental study [26]. We can use the model to explore longer contact times as long as timescales are shorter than those required to form mature focal adhesions. Given the lengths of time (less than 2 seconds or 4 seconds) from the experiment setup, including a mechanism for integrin clustering and maturity was deemed unnecessary and inaccurate. Furthermore, while the inclusion of a stochastic motor-clutch mechanism for the integrin dynamics is beneficial, given the computationally intensive task of fitting the parameters, this approach was deemed computationally unfeasible. Consequently, we utilised a piecewise linear function as an approximation for this mechanism. This mechanism was shown to be a suitable approximation in the research by Irons et al. [21]. The switching of the lighting during the cell’s contact time with the surface has not previously been investigated experimentally or computationally. Our simulations predict that periodically switching the lighting generates fluctuations in the maximum cell detachment force. However, given the noise level in this kind of experiment, it is not yet clear whether the predicted forces would be discernible; however, an emerging pattern is noticeable. Due to the high variability across cellular experiments, this model should be calibrated before the computational simulations are performed.

In summary, we have developed a model that is capable of accounting for light-induced changes in the force exerted by a biomaterial on a cell via integrins. This model predicts a measure of the number of bound and unbound integrins between the cell and a light responsive material surface under different experimental conditions. It has been calbirated with previously published experimental data. The strength of the model lies in its ability to predict the outcome of multiple experimental conditions in a short time. This was exemplified using instrumental parameters (contact time and retraction speed) and the material stimulation parameter (irradiation frequency) By exploring the effect of a range of parameter values on cell adesion force rapidly, the model can substantially aid in the selection of suitable experimental conditions and shortening of the time required to explore the parameter space. In addition, the model is able to predict experimentally inaccessible variables, including the relative proportion of bound and unbound integrins.

## Supporting information

Supplementary document for the main manuscript

## ACKNOWLEDGMENTS

The Engineering and Physical Sciences Research Council (EPSRC) supported the project’s funding via the Centre of Doctoral Training (CDT) in Transformative Pharmaceutical Technologies [EP/S023054/1].

## DATA AVAILABILITY

All data and code used for running experiments and model fitting are available on a GitHub repository at GitHub.

## Notes

https://github.com/mzelzer/Spring_Model_SCFM_DV

### Competing Interest Statement

The authors have declared no competing interest.

### Summary of Updates

The parameter table in the main document and the supplementary document.

