## Supplementary document for the main manuscript for "Model of integrin binding as a function of light-induced variations in forces exerted on cells via RGD-terminated, photoswitchable azobenzene surfaces"

#### Experimental Data Processing

Force curve processing was done using the JPK Instruments data processing software (Version 6.1.198). Data was smoothed, and the baseline offset and tilt were corrected by subtracting a linear fit from the retract segment after the last rupture event. Next, the x-offset was adjusted so that the first crossing of the y-axis of the retract segment was at x=0. Finally, the cantilever bending is subtracted from the piezo height to get the vertical tip position.

#### Numerical Methods for Simulation

The equations required to generate a simulated value for force ($F$) were written in C++, and the numerical methods implemented were verified against results from [1, 2]. The numerical method selected for solving the differential equations was a 5th-order Adams-Bashforth method [3]. The step size ($h \approx 1 \times 10^{-5}$) was chosen based on an analysis of the stability of solutions from the numerical solvers, and the step size remained consistent for all simulation time lengths.

#### Parameter Estimation

The parameters estimated through numerical minimisation are shown in Table S2. The parameters selected for numerical estimation primarily focus on binding and unbinding the integrins between the cell and the biomaterial (layer $L_4$).

Selection for the initial parameters fed into the algorithm is chosen using the Latin Hypersquare (LHS) method ([4]), which was used from the `scipy.stats.qmc`, [5]. The reason for selecting the LHS method is to sample the entirety of the multivariate parameter space while minimising computational expense [6, 7]. The range of parameters inputted into the LHS is selected based on preliminary simulations providing low $||L||_2$ values and using these as possible regions for initial estimations.

The minimisation between the simulated and experimental force was performed using the **Nelder-Mead** algorithm ([8]) from the Python package `scipy.optimize.minimize`, [5]. Testing different algorithms included in the `scipy.optimize.minimize` concluded that the **Nelder-Mead** algorithm provided a quantitatively appropriate fit with realistic parameters. Another benefit of this algorithm is bounds placed on the parameters generated between physically relevant upper and lower limits, $p_i \in (0, 100]$. The restriction in bounds prevents the algorithm from converging on negative values or overly large parameter estimates.

As each minimisation is computationally costly, we relied on an averaged SCFS retraction curves rather than minimising each data curve separately. As the experimental time step was larger than $h = 10^{-5}$, simulated time series were down-sampled for the fit. The full output from the parameter estimation can be found on the GitHub. The files containing the parameter sets generated by the Latin Hypersquare method can be found under the name `Spring_Model_Parameter_Analysis.csv`. The file containing the optimised parameter sets can be found in `init_Param_all_OFF_LHS_Powell_L2_50.txt` and `init_Param_all_ON_LHS_Nelder_L2.txt`.

#### *Biological Relevance of Estimated Parameters*

The values in Table S1 were chosen using a combination of criteria, including the availability of estimates or experimental values in the literature, analysis of plausibility, and parameter estimation.

### Increasing Fitting Accuracy

Using the parameters listed in Table S1 and performing the fitting described in the methods section, the parameter estimates shown in Table S2 are obtained. These parameters provide a sufficient fit and a reasonable estimation for some of the datasets, including the 1-second contact time ($T_p$) in both the **OFF** and **ON** lighting and the prediction for the 3 seconds contact time in the **OFF** lighting.

Although the present model has limitations in its predictive strength, we were also interested in understanding if a broader parameter space would result in better predictions. The data generated during fitting with parameters from Table S1 shows that the possible parameters provide a suitably low $||L||_2$ value have a range. With the uncertainty introduced by the necessity to estimate a number of parameters from literature values, we thought exploring the model fit with an adjusted parameter set was justified.

We selected to change the parameters to be $k_3 = 5$ N/m, and $k_1 = 1$ N/m, to bring the `nelder-mead` algorithm to converge to a parameter set that is within the parameter bounds. Furthermore, the parameters fitted to the 1 second contact time data in the **OFF** state are $\rho_u$, $\alpha$, $\beta$, $\bar{\eta}$, $\hat{\eta}$, $\bar{\theta}$, and $\hat{\theta}$. The bounds are decided that binding rate parameters $0 \leq \bar{\eta}, \hat{\eta}, \bar{\theta}, \hat{\theta} \leq 2$, $\alpha$ and $\beta$ have the same magnitude. Although these parameters ensure the algorithm converges, they are not biologically suitable and have not been incorporated in the main predictions. Further parameter estimation for $\varepsilon_c$ and $\varepsilon_e$ are carried out on the data relating to the **ON** lighting. This gives the parameter values in Table S2, producing the simulated retraction curves in Figure S1.

**Table S1.** Table containing parameters included in the model and their biological description.

| Parameter | Value (unit) | Biological Description | Literature Source |
|---|---|---|---|
| $k_1$ | 0.1 ($Nm^{-1}$) | Spring constant for the Colavinacin A | For convergence |
| $k_3$ | 0.05 ($Nm^{-1}$) | Spring constant for the cell | 0.1-50 $pN(nm)^{-1}$, [9], [10], [11] |
| $\rho_u$ | Estimated ($Nm^{-1}$) | Spring constant for the integrins between the cell and the SCFS tip | Numerically Estimated |
| $\rho_l$ | 1.0 ($Nm^{-1}$) | Spring constant for the integrins between the cell and the c(RGDfK) terminated azobenzene surface | See values $\alpha$ and $\beta$ |
| $\alpha$ | Estimated | Scaling factor on the force from the integrins bound to the contracted c(RGDfK) terminated azobenzene molecules | Numerically Estimated |
| $\beta$ | Estimated | Scaling factor on the force from the integrins bound to the extended c(RGDfK) terminated azobenzene molecules | Numerically Estimated |
| $\bar{\eta}$ | Estimated | Rate of binding between the cell and the c(RGDfK) terminated azobenzene molecules in the contracted state | Numerically Estimated |
| $\hat{\eta}$ | Estimated | Rate of unbinding between the cell and the c(RGDfK) terminated azobenzene molecules in the contracted state | Numerically Estimated |
| $\bar{\theta}$ | Estimated | Rate of binding between the cell and the c(RGDfK) terminated azobenzene molecules in the extended state | Numerically Estimated |
| $\hat{\theta}$ | Estimated | Rate of unbinding between the cell and the c(RGDfK) terminated azobenzene molecules in the extended state | Numerically Estimated |
| $\varepsilon_c$ | Estimated | Rate of un/binding when the c(RGDfK) terminated azobenzene surface is in the **ON** state | Numerically Estimated |
| $\varepsilon_e$ | Estimated | Rate of un/binding when the c(RGDfK) terminated azobenzene surface is in the **OFF** state | Numerically Estimated |
| $g_1$ | 0.02 ($s^{-1}$) | Unstressed binding rate | [12], [13] |
| $h$ | 10.0 (nm) | Integrin binding | [14] |
| $s$ | 15.0 (nm) | Maximum integrin binding range | [14] |
| $l_1$ | 0.005 ($s^{-1}$) | Unstressed unbinding rate between cell and c(RGDfK) terminated azobenzene surface | [12], [15] |
| $l_2$ | 0.01 ($s^{-1}$) | unbinding parameter between the cell and the c(RGDfK) terminated azobenzene surface | Model choice |
| $l_3$ | 70.0 ($s^{-1}$) | forced unbinding rate between the cell and the c(RGDfK) terminated azobenzene surface | Model choice |
| $u_1$ | 0.02 ($s^{-1}$) | Unstressed unbinding rate between cell and the Concanavalin A | [12], [13] |
| $u_2$ | 0.01 ($s^{-1}$) | unbinding rate parameter between the cell and the ECM | Model choice |
| $u_3$ | 1.0 ($s^{-1}$) | forced unbinding rate between the cell and the ECM | Model choice |
| $h$ | 10 (nm) | Maximum binding range | [14] |
| $s$ | 15 (nm) | Maximum unbinding range | [14] |

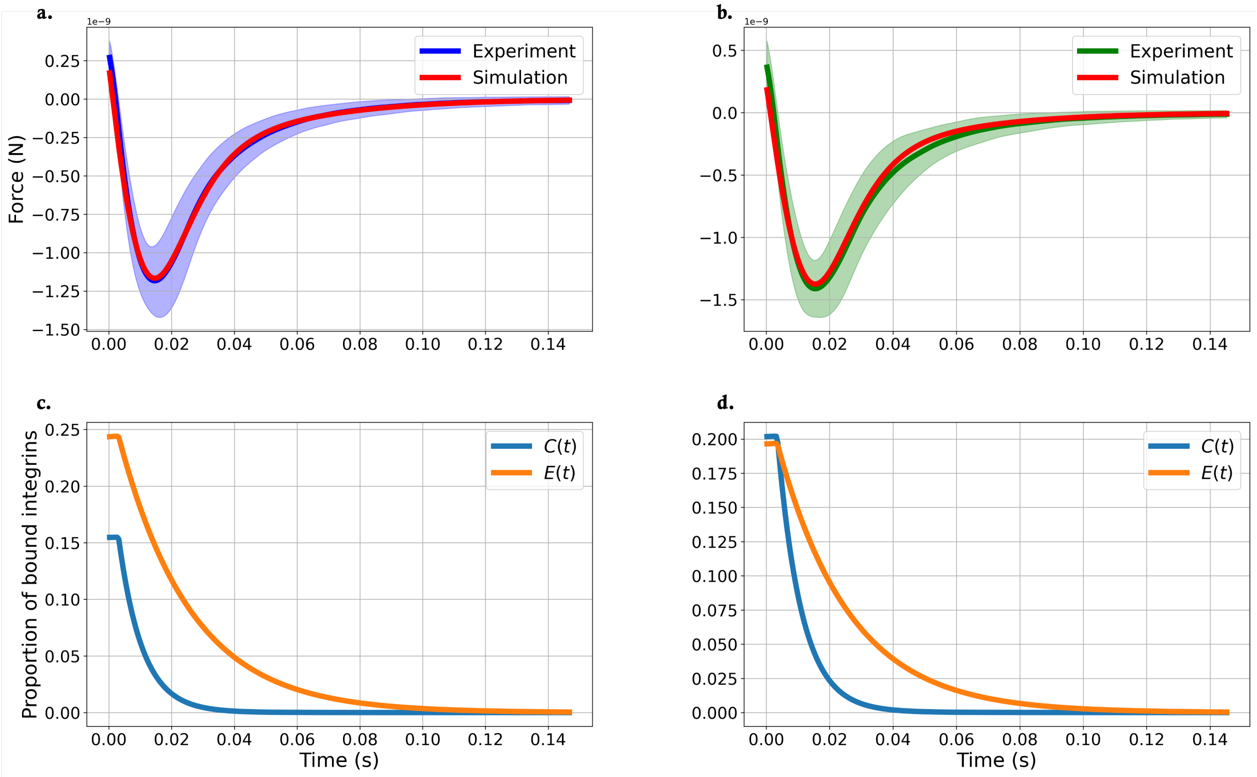


**Figure S1.** Simulated and average experimental force retraction curves in the **OFF** (a) and **ON** (b) states and simulated relative integrin binding in both states (c, d). In (a) and (b), experimental data (dark purple and dark green lines) is shown with one standard deviation ($\pm\sigma$, light purple and light green areas). Simulated retraction curves (red lines) were generated using the optimised parameter set (Table S2). (c) and (d): Total relative proportion ($C$, $E$) of integrin binding to c(RGDfK) terminated azobenzene surfaces in the **OFF** (c) and **ON** (d) state as the SCFS probe is retracted over time. Integrin binding is a relative proportion of the total number of bound integrins.

| $\|L\|_2$:**OFF** | $\|L\|_2$:**ON** | $\rho_u$ nN(nm)$^{-1}$ | $\alpha$ | $\beta$ | $\bar{\eta}$ | $\hat{\eta}$ | $\bar{\theta}$ | $\hat{\theta}$ | $\varepsilon_c$ | $\varepsilon_e$ |
|---|---|---|---|---|---|---|---|---|---|---|
| 0.374 | 0.592 | 0.00871 | 0.129 | 0.0033 | 1.308 | 1.245 | 1.911 | 0.633 | 0.766 | 0.776 |

**Table S2.** Estimated parameters using $k_3 = 5$ nN(nm)$^{-1}$, and $k_1 = 1$ nN(nm)$^{-1}$.

The simulated retraction curves obtained with these higher values for $k_1$ and $k_3$ (Figure S1) align more closely with the averaged retraction curve. In addition, the variation in $F_{\max}$ as a function of varying $T_p$ (Figure S2) becomes more pronounced between the 1-second and 3-second contact time data. This more closely reflects the experimental observations made in Kadem *et al.* [16], suggesting that these new values for $k_1$ and $k_3$ provide fits to the model that match the experimental data better. Given that the values used for $k_1$ and $k_3$ are not realistic based on existing mechanobiological data in the literature, the results in Figures S1 and S2 can only indicate the possibility of the model to provide better fits if the parameter space can be defined better with experimental data.

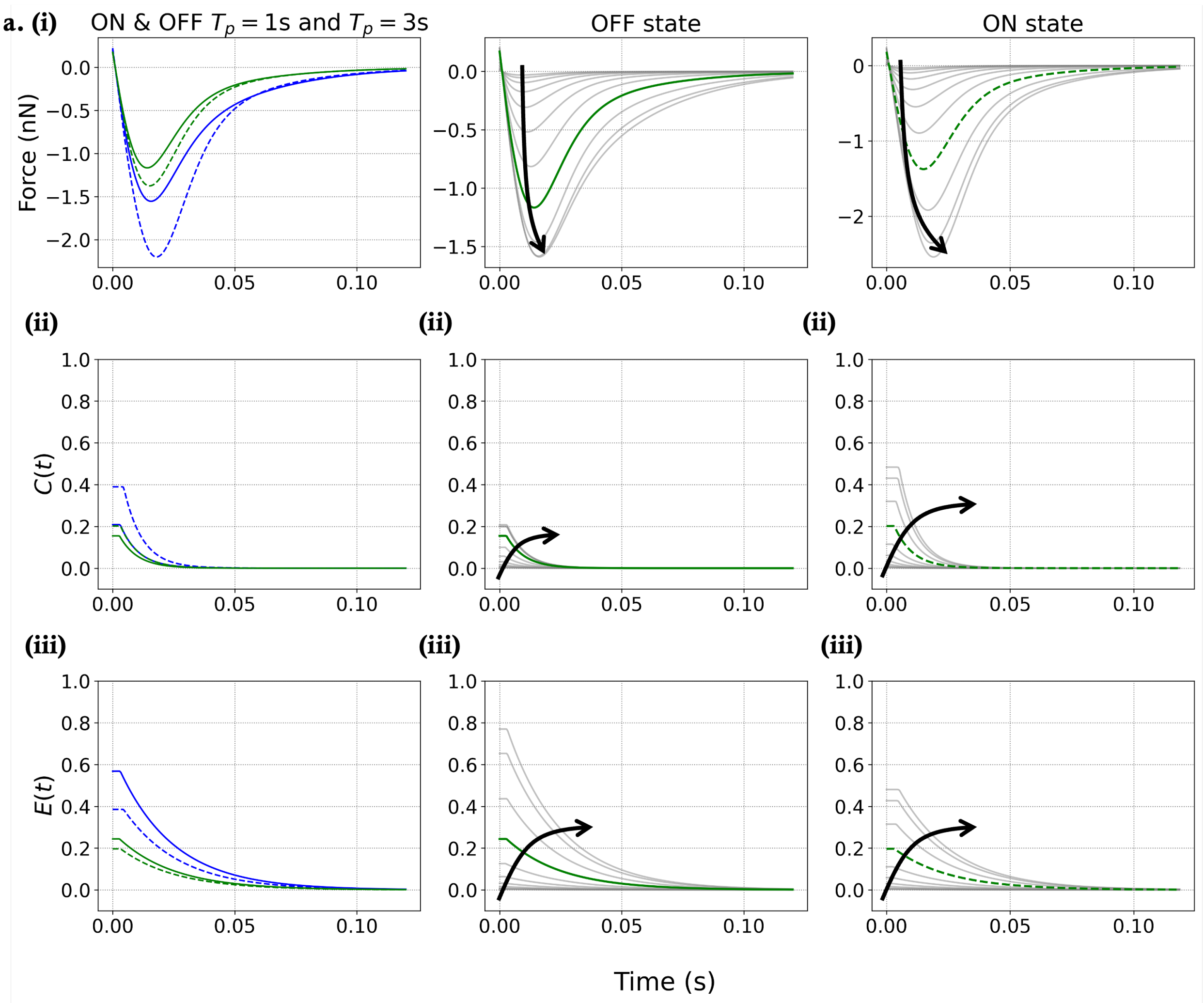


**Figure S2.** Simulated force curves (i) and relative amount of integrins bound to *cis*-azobenzene ($C$) (ii) and *trans*-azobenzene ($E$) (iii) for varying cell-surface contact times ($T_p$) on surfaces in either the **OFF** or **ON** state. (a) Simulated data for the **OFF** (solid lines) and **ON** (dotted lines) state for $T_p = 1$ s (green) and $T_p = 3$ s (blue). (b,c) Simulated data for a range of contact times ($T_p \gg 1$ and $T_p \ll 1$) for surfaces in either the **OFF** (b) or **ON** (c) state. The (green) line represents $T_p = 1$ s; $\cdots$ indicates $T_p > 1$ s; $\cdot - \cdot$ indicates $T_p < 1$ s. For all parameters other than $T_p$, the optimised and predefined parameter values given in Tables S2 and S1 are used. For the **ON** state, $\lambda_c = \varepsilon_c$ and $\lambda_e = \varepsilon_e$. For the **OFF** state, $\lambda_c = 0.2$ and $\lambda_e = 0.8$.

[5] Pauli Virtanen, Ralf Gommers, Travis E. Oliphant, Matt Haberland, Tyler Reddy, David Cournapeau, Evgeni Burovski, Pearu Peterson, Warren Weckesser, Jonathan Bright, Stéfan J. van der Walt, Matthew Brett, Joshua Wilson, K. Jarrod Millman, Nikolay Mayorov, Andrew R. J. Nelson, Eric Jones, Robert Kern, Eric Larson, C J Carey, İlhan Polat, Yu Feng, Eric W. Moore, Jake VanderPlas, Denis Laxalde, Josef Perktold, Robert Cimrman, Ian Henriksen, E. A. Quintero, Charles R. Harris, Anne M. Archibald, Antônio H. Ribeiro, Fabian Pedregosa, Paul van Mulbregt, and SciPy 1.0 Contributors. SciPy 1.0: Fundamental Algorithms for Scientific Computing in Python. *Nature Methods*, 17:261–272, 2020.
